# Genome-wide Discovery of lncRNAs in Mucorales Reveals Essential Roles in Development and Fungal Biology

**DOI:** 10.1101/2025.10.01.679568

**Authors:** Ghizlane Tahiri, Hrant Hovhannisyan, Carlos Lax, Eusebio Navarro, Toni Gabaldón, Francisco E. Nicolás, Victoriano Garre

## Abstract

Long non-coding RNAs (lncRNAs) emerged as key regulators across eukaryotes, yet their functions in early-diverging fungal (EDF) pathogens remain largely unknown. Here, we provide the first comprehensive identification and characterization of lncRNAs in the EDF order Mucorales, a threatening and WHO high-priority group of opportunistic human pathogens. In this work, we focus on the two major models of this group: *Mucor lusitanicus* and the clinically relevant pathogen *Rhizopus microsporus*. We show that EDF lncRNAs exhibit conserved features, dynamic regulation during host interactions, and integration within critical gene regulatory networks. Despite lncRNAs being preferentially encoded in inactive chromatin compartments, we found that their expression is associated with 6mA presence in *R. microsporus*. Additionally, we also found that lncRNAs can be targeted by both the canonical RNA interference pathway and the non-canonical RNA interference mechanism. Comparative genomics revealed a subset of evolutionarily conserved lncRNAs, including two essential for fungal viability (lncRNA2 and lncRNA4). LncRNA4 disruption, even in heterokaryosis, resulted in severely affected growth and filamentation. These results establish lncRNAs as indispensable regulators of fungal physiology and pathogenicity, highlighting their potential as novel antifungal targets.

## INTRODUCTION

Defined as RNA transcripts longer than 200 nucleotides with limited or no protein-coding potential, lncRNAs influence numerous biological processes, including chromatin remodeling, transcriptional control, and post-transcriptional regulation^1^. They undergo canonical RNA polymerase II-dependent transcription and can be processed similarly to protein-coding genes, with 5’ capping, splicing, and polyadenylation^2^; non-polyadenylated forms are stabilized via alternative mechanisms, such as circularization or interactions with ribonucleoproteins^3^. LncRNAs exhibit low basal expression but high context-specificity, with tightly regulated spatial, temporal, and condition-dependent patterns^4^. Their subcellular localization further determines function: nuclear lncRNAs are often involved in transcriptional regulation and epigenetic modifications, whereas cytoplasmic lncRNAs influence mRNA stability, translation, or protein activity^5,6^. Despite rapid sequence evolution and generally low conservation, promoter regions and certain positional features are frequently maintained, suggesting functional constraints across divergent taxa^4^.

In fungi, lncRNAs have been identified as important regulators of development, metabolism, stress responses, and pathogenicity^7,8,9^. In *Saccharomyces cerevisiae*, several lncRNAs such as IRT1, RME2, and RME3 control meiotic gene expression by acting in cis or antisense to target genes, often through recruitment of chromatin-modifying complexes^2^. In *Schizosaccharomyces pombe*, lncRNAs such as the meiosis-specific meiRNA regulate gene expression and nuclear organization by forming nuclear foci (Mei2 dots) that sequester the silencing factor Mmi1, thereby stabilizing meiotic transcripts, facilitating homologous chromosome pairing, and influencing heterochromatin assembly and RNAi-mediated gene silencing^10,11^. Similarly, in pathogenic fungi, lncRNAs have been implicated in regulating virulence, morphogenesis, secondary metabolism, and stress responses^7^. For example, the nuclear lncRNA RZE1 in *Cryptococcus neoformans* modulates the yeast-to-hyphae switch, ncRNA1 in *Ustilago maydis* acts in cis to influence pathogenicity^12^, and genome-wide studies in *Candida* species reveal hundreds of lncRNAs co-expressed with protein-coding genes and differentially regulated during infection, suggesting roles in host-pathogen interactions^13^. Beyond pathogenic yeasts, lncRNAs have also been reported in filamentous fungi such as *Aspergillus* species, where they respond to environmental stress and may regulate metabolism and cellular survival under stress conditions^14^. However, these studies are largely restricted to Dikaryotic model organisms, leaving early-diverging lineages, including Mucorales, largely unexplored.

Some Mucorales representatives, including *Mucor lusitanicus* and *Rhizopus microsporus*, are opportunistic human pathogens causing mucormycosis, a rapidly progressing and often fatal infection that primarily affects immunocompromised individuals^15^. Yet, the molecular mechanisms underpinning gene regulation, development, and virulence in Mucorales, particularly those mediated by lncRNAs, remain poorly understood.

The study of lncRNAs in Mucorales offers a unique opportunity to gain insight into the evolutionary origin of RNA-based regulation and its role in fungal biology, sheding light on ancient regulatory mechanisms and their contributions to fungal adaptation and pathogenicity. In addition, lncRNAs playing essential or virulence-related roles could serve as therapeutic targets.

Here, we present a comprehensive analysis of lncRNAs in *M. lusitanicus* and *R. microsporus*, combining deep RNA sequencing across developmental stages, stress conditions, macrophage exposure, and murine infection. Using a stringent computational pipeline integrating transcript assembly, coding potential assessment, and genomic context, we identified hundreds of high-confidence lncRNAs per species. These lncRNAs display canonical features of non-coding transcripts, dynamic expression patterns, and positional enrichment near genes involved in virulence, stress response, and metabolism. Comparative analyses reveal limited sequence conservation but highlight conserved regulatory features, supporting the notion of lineage-specific lncRNA evolution with occasional evolutionary persistence.

Collectively, these findings uncover a previously unrecognized layer of transcriptional complexity in Mucorales, establishing lncRNAs as integral components of fungal regulatory networks. Our study provides a foundation for future functional investigations and suggests that lncRNA-mediated control is a conserved and adaptable feature of early diverging fungal lineages, with implications for fungal pathogenesis and potential antifungal strategies

## MATERIAL AND METHODS

### RNA-seq Datasets processing

To identify long non-coding RNAs (lncRNAs) in *Mucor lusitanicus* and *Rhizopus microsporus*, we utilized publicly available RNA-seq datasets derived from in vitro and in vivo interactions of fungal spores with macrophages and mice infection. Public datasets were retrieved from the Sequence Read Archive (SRA) database (16) (last access 3^rd^ August 2022) using SRA Toolkit v2.9.6.1 with the prefetch and fastq-dump commands. In total, 44 public RNA-seq libraries were analyzed for *M. lusitanicus* (17,18,19) and 17 datasets for *R. microsporus* (20,21).

Raw reads underwent quality assessment with FASTQC v0.11.8 (https://www.bioinformatics.babraham.ac.uk/projects/fastqc/) and MultiQC v1.0 (22), followed by adapter trimming and removal of low-quality bases using Trimmomatic v0.36 (23) with the following parameters: ILLUMINACLIP:TruSeq3_adapters.fa:2:30:10 LEADING:3 TRAILING:3 SLIDINGWINDOW:4:15 MINLEN:49. Cleaned reads were aligned to the respective reference genomes obtained from the the Joint Genome Institute (JGI) MycoCosm database (accessed August 3, 2022) (24) using TopHat2 v2.1.1 (25) with the –b2-very-sensitive and –max-intron-length 1000 options, reflecting typical fungal intron sizes. For dual RNA-seq datasets containing both fungal and mouse transcripts, reads were mapped to concatenated and mouse reference genomes (mouse genome from Ensembl release 107, accessed January 30, 2023) (http://www.ensembl.org/Mus_musculus/Info/Index) (26), followed by extraction of fungus-specific alignments using Samtools v1.3.1 (27).

### Transcriptome assembly and lncRNA prediction

Genome-guided transcriptome assemblies were generated for each sample BAM file with StringTie v1.3.4b (28). Resulting GTF files were merged per species using StringTie merge (–g 50). Novel transcripts longer than 200 bp were identified by comparing merged assemblies to reference annotations using Gffcompare v0.11.2 (https://ccb.jhu.edu/software/stringtie/gffcompare.shtml), retaining only intergenic transcripts for further analysis.

Coding potential was assessed using two complementary tools: CPC v0.9 (29), run against the UniProt database, and FEELnc v0.1.1 (30) employing a random forest classifier trained with protein-coding sequences in shuffle mode. Transcripts classified as non-coding by both tools were retained as putative lncRNAs. Transcripts flagged by FEELnc for ambiguous nucleotides but confirmed non-coding by CPC were also included. For global expression analyses, gene and lncRNA expression levels were quantified using FeatureCounts v1.6.4 (31). Counts were normalized to transcripts per million (TPM) to account for transcript length and library size.

### Integrative analysis of lncRNA with epigenetic data

lncRNA density in 20 kb bins was computed and compared with gene and TE density, as well as 6mA, H3K4me3, H2A.Z, and H3K9me3 genome-wide distribution retrieved from our previous work (21). Spearman correlation was computed between each feature using deepTools (32) MultiBigWig summary tool v3.1. 6mA profile graphs over lncRNAs and genes were generated by computing 6mA frequency over equal-size bins using deepTools v3.1 (32). Genes with TPM = 0 were considered as silent, whereas genes with TPM > 0 were considered as transcriptionally active. MACs absence/presence was computed within ± 300 bp from the TSS. To visualize the effects of MAC loss in gene expression, we used the Integrative Genome Viewer (IGV) (33).

To address the implications of RNAi machinery in lncRNA regulation, sRNA and mRNA reads from WT and *r3b2*Δ strains, with and without macrophages (19), were processed as follows. Raw sRNA reads were quality-checked with FASTQC before and after adapter removal. Adapter sequences, contaminant reads, and low-quality reads (-q28 - p50) were removed using TrimGalore! v0.6.2 (https://github.com/FelixKrueger/TrimGalore). Clean sRNA reads (18–35 nt) were aligned to the *M. lusitanicus* v2.0 genome using Bowtie2 v2.5.3 (34). Mapped reads were quantified with FeatureCounts v1.6.4. Raw mRNA reads were aligned to the same genome using HISAT2, quantified with FeatureCounts v1.6.4, and differential expression analysis for both sRNA and mRNA was conducted with DESeq2 v1.30.0 (35), considering transcripts with FDR ≤0.05 and |log₂FC| ≥1. Scatter plots were generated to examine the correspondence between sRNA production and lncRNA expression.

### Phylogenetic tree construction and evolutionary conservation analyses

Proteomes of *M. lusitanicus*, *R. microsporus*, *R. microsporus* var. *chinensis*, *Amylomyces rouxii*, *R. delemar*, *R. stolonifer*, *Sporodiniella umbellata*, *Ellisomyces anomalus*, *Mucor ambiguus*, *M. circinelloides*, *Parasitella parasitica*, and *M. racemosus* were retrieved from the Joint Genome Institute (JGI) MycoCosm genome portal (24). Orthogroups were inferred from all-versus-all sequence similarity searches using OrthoFinder (v2.5.5) (36), and single-copy orthologs present in the majority of species were aligned using MAFFT (v7.471) (37). The resulting alignments were concatenated, trimmed, and used to infer a species tree with RAxML (raxmlGUI v2.0.10) (38,39) under the PROTGAMMAWAGF substitution model with 1000 bootstrap replicates. The final phylogeny was visualized and annotated using the Interactive Tree of Life (iTOL, v6) (40). To assess lncRNA conservation across Mucorales species, genomes of the selected species were compiled into BLAST databases, and predicted lncRNAs were queried using BLASTn BLASTn v2.9.0 (41) (allowing up to five target sequences per query, a maximum of five high-scoring segment pairs, and an e-value threshold of 1e-3). Hits covering at least 50% of the query length were retained and converted into BED files, which were then intersected with annotated genomic features, including tRNAs and protein-coding genes (PCGs) using bedtools v2.29.2 (https://bedtools.readthedocs.io/en/latest/). LncRNAs that aligned to tRNAs or PCGs were discarded from the conservation analyses, and only those mapping to unannotated regions were retained, enabling the identification of conserved and species-specific lncRNAs

### Fungal strains and culture conditions

All the *R. microsporus* strains used in this work derived from the WT *Rhizopus microsporus* ATCC 11559 (listed in Supplementary Table 1). The avirulent control was the UM33, which was employed as the receptor strain for the mutant generation. All *R. microsporus* strains were cultured in rich medium YPG pH 4.5 at 30°C for spore harvesting, supplemented with uridine (200 mg/l) when required. All the media and culture conditions used *R. microsporus* growth are detailed in Lax et al. (42).

### Mutant strain generation by CRISPR-Cas9

*R. microsporus* mutant strains were generated following previously established procedures (42,43). Three infection-induced lncRNAs (lncRNA1: MSTRG.4436, lncRNA2: MSTRG.1050, and lncRNA3: MSTRG.7756) and one conserved lncRNA (lncRNA4: MSTRG.4826) were selected for targeted disruption using the CRISPR-Cas9 system. Guide RNAs (crRNAs) were designed to target the first exon of each lncRNA with EuPaGDT (http://grna.ctegd.uga.edu/) under default parameters (Supplementary Table 2). For homology-directed repair, the *pyrF* selectable marker was PCR-amplified with 38 bp homology arms flanking the Cas9 cut site. PCR amplifications were carried out using Herculase II fusion DNA polymerase (Agilent), and fragments were gel-purified when necessary. Primers (Supplementary Table 3) were designed with Primer3Stat (http://bioinformatics.org/sms2/pcr_primer_stats.html) and the Multiple Primer Analyzer (Thermo Fisher). Protoplasts of the auxotrophic UM33 strain (*pyrF-*, *LeuA-*) were transformed by electroporation with Cas9–gRNA ribonucleoprotein complexes and linear repair templates (150–200 ng/μl). Transformants were selected on minimal media lacking uridine (MMC selective medium). Integration of *pyrF* and homokaryosis were confirmed by diagnostic PCR using primers targeting *pyrF* and the flanking genomic regions, following the procedure described by Lax et al (42).

### Phenotypic characterization of the lncRNA mutants

Radial growth and sporulation of the phagocytosis mutants were analysed, inoculating drops of 500 spores in the plate center of MMC 3.2 (for radial growth) and YPG 4.5 (for sporulation). For each strain, three independent plates were prepared, and three measurements per colony were taken daily. Colony radial growth was assessed over 5 days, with measurements of colony diameter taken every 24 hours. Radial growth was quantified using three replicates per mutant. Statistical analysis was conducted using one-way ANOVA followed by Student’s *t*-test, considering differences statistically significant at *p* ≤ 0.05.

### RNA extraction, library preparation, and RNA-seq analysis of lncRNA mutants

Three replicates of the fungal spores from the WT strain (UM6) and the lncRNA mutants (lncRNA2 and lncRNA4) growing in saprophytic conditions in MMC media, under light and 30°C, for 24 hours. Total RNA was extracted using the RNeasy Mini Kit (Qiagen, Hilden, Germany) following the manufacturer’s instructions. RNA library preparation was carried out by a commercial sequencing provider. Plate-based RNA sample prep was performed using the PerkinElmer Sciclone NGS robotic liquid handling system with Illumina’s TruSeq Stranded mRNA HT sample preparation kit, incorporating poly(A) selection. Libraries were prepared using 1 μg of total RNA per sample and amplified with 8 PCR cycles, following the Illumina user guide (support.illumina.com/sequencing/sequencing_kits/truseq-stranded-mrna.html). Libraries were quantified using KAPA Biosystems’ qPCR library quantification kit (Roche) on a LightCycler 480 instrument and pooled for multiplexing. Sequencing was performed on an Illumina NovaSeq 6000 platform using NovaSeq XP v1.5 reagent kits (S4 flow cell) with a 2 × 150 bp paired-end configuration.

Raw read datasets were processed following the same procedure described above for *M. lusitanicus*, using the *R. microsporus* v2.0 genome (version 2.0; https://mycocosm.jgi.doe.gov/Rhimi59_2/Rhimi59_2.home.html) (24). Differential expression analysis was performed with DESeq2 (FDR ≤0.05, |log₂FC| ≥1), and all differentially expressed genes and lncRNAs from all conditions are provided in Supplementary Data 4.

### Co-expression network construction and module–trait association analysis

Normalized expression matrices from *R. microsporus* wild-type spores grown under saprophytic conditions (n = 3) and during host–pathogen interaction (n = 3) were used for network analysis. Low-variance and low-prevalence genes were filtered prior to analysis. Weighted gene co-expression network analysis (WGCNA) was performed to identify co-expression modules from differentially expressed protein-coding genes (PCGs) and lncRNAs. A soft-thresholding power of β = 50 was chosen based on the scale-free topology criterion (R² > 0.8), with parameters deepSplit = 2, mergeCutHeight = 0.3, and minModuleSize = 30. Module eigengenes (MEs) were correlated with infection status to identify modules significantly associated with the trait (adjusted p-value < 0.05). Modules containing lncRNAs of interest were prioritized, and genes within modules were ranked by module membership (MM > 0.6) and gene significance (GS > 0.6).

### Gene density distribution

Scaffold sizes were extracted from the *R. microsporus* reference genome to ensure consistency with the genomic structure. Fixed 50 kb windows were generated across the genome and aligned with gene annotation data. Gene density was calculated by mapping gene lengths onto each 50 kb window, and the results were exported as BEDGraph files. Circos (44) was used to visualize genome-wide gene density and the distribution of upregulated and downregulated protein-coding genes (PCGs) and lncRNAs in the lncRNA4⁻ mutant.

## RESULTS

### Identification of intergenic lncRNAs in Mucorales

To explore the landscape of lncRNAs in Mucorales, we analyzed high-throughput RNA sequencing (RNA-seq) data from two representative species: *Mucor lusitanicus* and *Rhizopus microsporus* (**Figure 1**). We compiled a total of 44 RNA-seq datasets from *M. lusitanicus*, encompassing multiple infection stages in confrontation assays with macrophages, the primary immune cells involved in fungal clearance. For *R. microsporus*, we used 17 RNA-seq datasets derived from an *in vivo* murine mucormycosis model and different environmental conditions. These previously published RNA-seq datasets were processed for quality control, adapter trimming, and removal of low-quality reads.

**Fig. 1.**
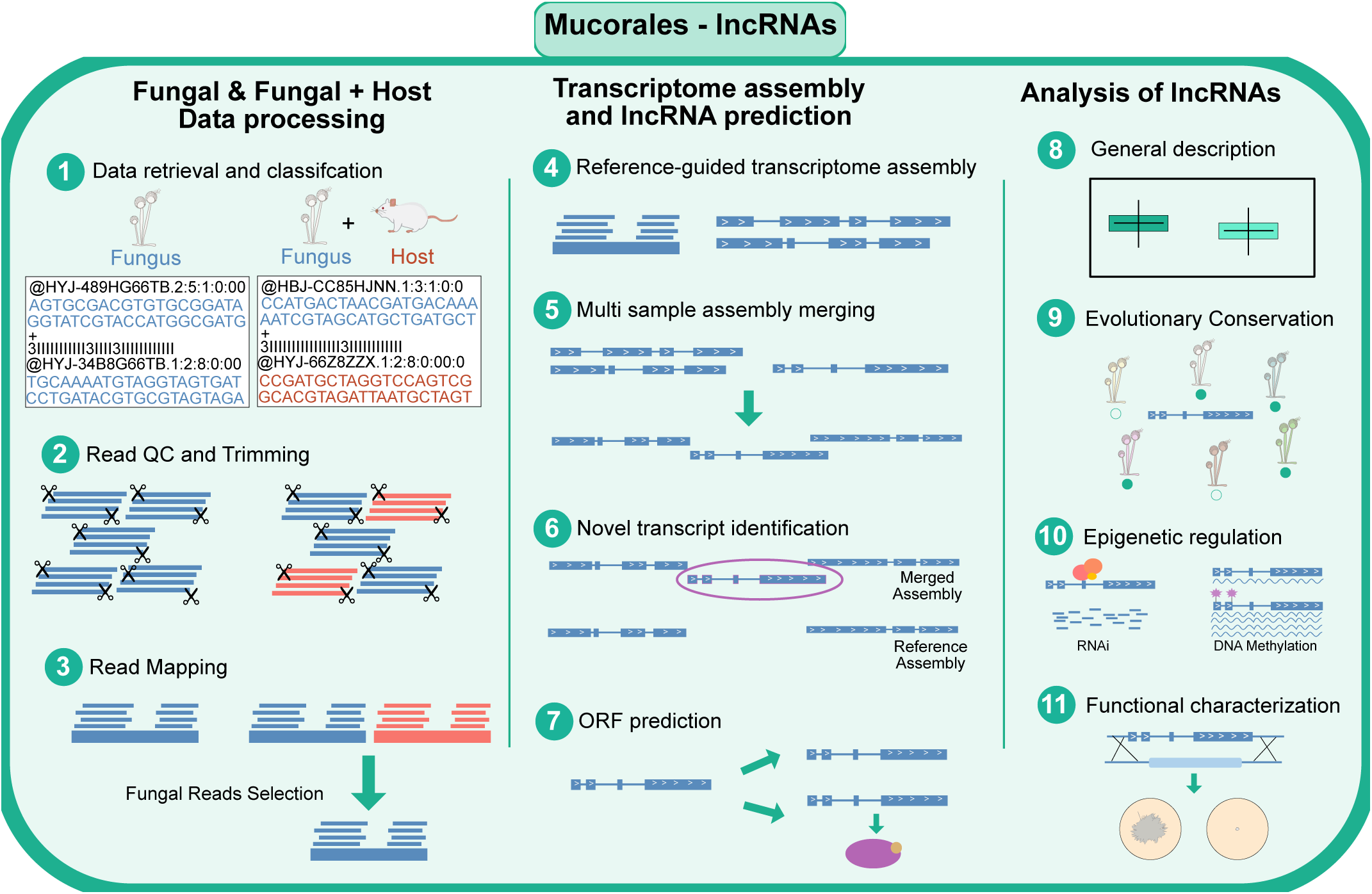
Pipeline for the identification and characterization of lncRNAs in Mucorales. Schematic representation of the bioinformatics workflow used to identify long non-coding RNAs (lncRNAs) in *Mucor lusitanicus* and *Rhizopus microsporus*. RNA-seq libraries from infection and control conditions were subjected to quality control, adapter trimming, and alignment to fungal and host genomes. Fungal reads were extracted and used for genome-guided transcriptome assembly with StringTie, followed by transcript merging. Novel intergenic transcripts longer than 200 nucleotides were selected. Coding potential was assessed using CPC and FEELnc, retaining only transcripts predicted as non-coding by both tools. Final lncRNA candidates were analyzed for expression, genomic distribution, conservation, epigenetic regulation, and functional characterization.

Transcriptome assemblies were generated for each sample and species, and individual assemblies were subsequently merged to create a non-redundant reference transcriptome for each species. Novel transcripts not overlapping with existing annotations were identified and retained for downstream analysis (**Figure 1**). To classify transcripts as lncRNAs, two main criteria were applied: (i) transcript length greater than 200 nucleotides, and (ii) lack of protein-coding potential. Coding potential was evaluated using the Coding Potential Calculator (CPC), which assesses the features of open reading frames and sequence similarity to known protein sequences, and FEELnc, which employs a random forest classifier trained on protein-coding sequences. Only transcripts classified as non-coding by both methods were retained as putative lncRNAs (**Figure 1**). Due to the non-strand-specific nature of the utilized RNA-seq libraries, only intergenic lncRNAs, transcripts located in genomic regions devoid of annotated protein-coding genes, were considered. This class of lncRNAs is also the most prevalent in other fungal systems^13,16^. This pipeline allowed us to identify 622 intergenic long non-coding RNAs (lncRNAs) in *Mucor lusitanicus* and 470 in *Rhizopus microsporus* (**Supplementary Data 1**) (**Figure 2A**).

**Fig. 2.**
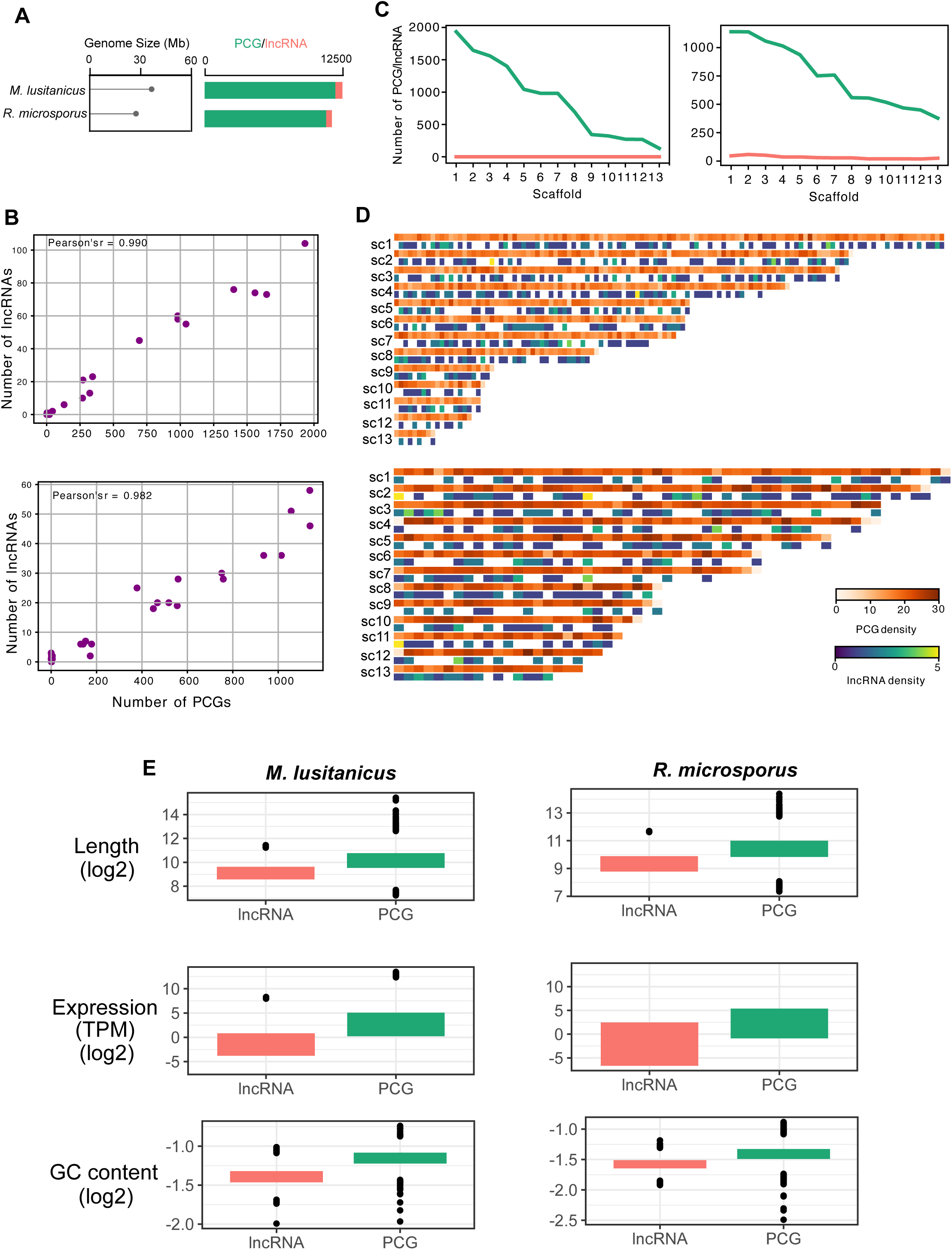
Genomic landscape of lncRNAs and protein-coding genes in *M. lusitanicus* and *R. microsporus*. (**A**) Genome size, number of protein-coding genes (PCGs), and number of identified intergenic lncRNAs. *M. lusitanicus* genome: 36.6 Mbp; *R. microsporus* genome: 27.39 Mbp. **(B)** Distribution of lncRNAs and PCGs per scaffold (macro-scale). Lines represent the total number of genes per scaffold for each type. **(C)** Correlation between lncRNA and PCG counts per scaffold. Scatter plots show strong positive correlations (Pearson’s r = 0.990 for *M. lusitanicus*, r = 0.982 for *R. microsporus*). **(D)** Distribution of gene density across scaffolds segmented into 50 kb bins (micro-scale). Each row represents a scaffold labeled by gene type (PCG or lncRNA); colors indicate gene counts per bin. lncRNAs and PCGs frequently co-localize in gene-dense regions. **(E)** Comparison of expression levels, transcript lengths, and GC content between lncRNAs and PCGs. Boxplots show significant differences (Wilcoxon rank-sum test, P < 0.05); lncRNAs have lower expression, shorter transcripts, and reduced GC content.

### Genomic distribution and sequence characteristics of lncRNAs in Mucorales

To explore the genomic distribution of the identified lncRNAs in *M. lusitanicus* and *R. microsporus*, we first mapped their positions across assembled scaffolds. The number of lncRNAs per scaffold showed a strong positive correlation with the number of PCGs (Pearson’s r = 0.990 in *M. lusitanicus*, r = 0.982 in *R. microsporus*; **Figure 2B**), which, given their intergenic nature, resulted in a higher lncRNA density scaled with gene PCG abundance (**Figure 2B, 2C, and Supplementary** Figure 1). PCG and lncRNA density calculation in 50 kb bins also showed a high frequency of colocalization of these two elements in both species (**Figure 2D**). In addition,, expression levels, GC content, and transcript length of lncRNAs were analyzed (**Figure 2E**). In *M. lusitanicus*, lncRNAs exhibited significantly lower expression than PCGs under both macrophage confrontation conditions and non-stressful growth, and a similar trend was observed in *R. microsporus* (**Figure 2E**). These findings are consistent with previous reports across a broad range of eukaryotic systems, where lncRNAs are typically expressed at lower levels than coding genes^5,9,17^. Mucorales lncRNAs also displayed significantly lower GC content compared to PCGs (**Figure 2E**), suggesting conserved structural or regulatory constraints distinct from those acting on coding sequences. This difference aligns with observations in other fungi^9,17,18^. Trancript lengths also varied between PCGs and lncRNAs, with the latter showing significantly shorter length in both species. Identified lncRNAs in these early diverging branches of fungal evolution suggest that they may be subject to evolutionary and functional constraints distinct from those of protein-coding genes across the tree of life.

### lncRNAs are preferentially distributed in inactive chromatin regions and their expression is associated with 6mA

EDF are characterized by having a distinct epigenetic landscape compared to most eukaryotes. Instead of 5-methylcytosine (5mC) being the main DNA epigenetic mark, several of these organisms rely on N6-methyladenine (6mA) as the principal epigenetic regulator of DNA, a feature shared with bacteria^19^. We previously characterized that this epigenetic modification is associated with the activation of gene expression in Mucorales and, in the case of *R. microsporus*, its presence is essential for survival^20,21^. This fungus exhibits a compartmentalized chromatin landscape, with active regions enriched in 6mA and H3K4me3, and inactive regions where these modifications are less frequent but transposons and H3K9me3 predominate^21^. Genomic locations of lncRNA genes showed a stronger positive correlation with inactive chromatin regions (**Figure 3A**), where activation-associated marks are less abundant. This may explain the generally lower expression levels of lncRNAs as compared to PCGs. Nonetheless, given the essential role of 6mA in regulating gene expression, we analyzed the distribution of this modification across lncRNA genes. We found that although its frequency was lower than in PCGs, its distribution pattern was similar, with 6mA marks mainly concentrated downstream of the TSS and within promoter regions (**Figure 3B**). This prompted us to investigate whether the presence of 6mA was differentially associated with silent (TPM = 0) or transcriptionally active lncRNA genes. Similar to observations made for PCGs^20,21^, 6mA was more frequent in active than in silent lncRNA genes (96 and 374 lncRNAs, respectively) (**Figure 3C**). In the *R. microsporus* genome, this modification typically appears in specific regions termed methylated adenine clusters (MACs)^21^. Consistently, the expression of lncRNA genes harboring a MAC around the TSS (−300 to +300 bp) was higher than that of those lacking one. Indeed, methylated lncRNA genes (MAC present) exhibited higher expression levels compared to unmethylated ones (without MAC) (p-value = 0.0059) (**Figure 3D**). This association led us to explore the dynamics of 6mA changes and their impact on lncRNA expression. In previous work, we developed a knockdown of the main 6mA DNA methyltransferase in *R. microsporus*, Mta1, using ZnSO₄^21^. We examined lncRNA loci that lost a MAC due to reduced Mta1 activity and found that 12 out of 17 differentially expressed lncRNAs were repressed, as exemplified by MSTRG.5861 (**Figure 3E and Supplementary Data 2**), while 5 were upregulated. Taken together, these results show that although lncRNA genes are preferentially distributed in inactive chromatin regions, 6mA is nevertheless clearly associated with their expression, paralleling the regulation observed in PCGs.

**Fig. 3.**
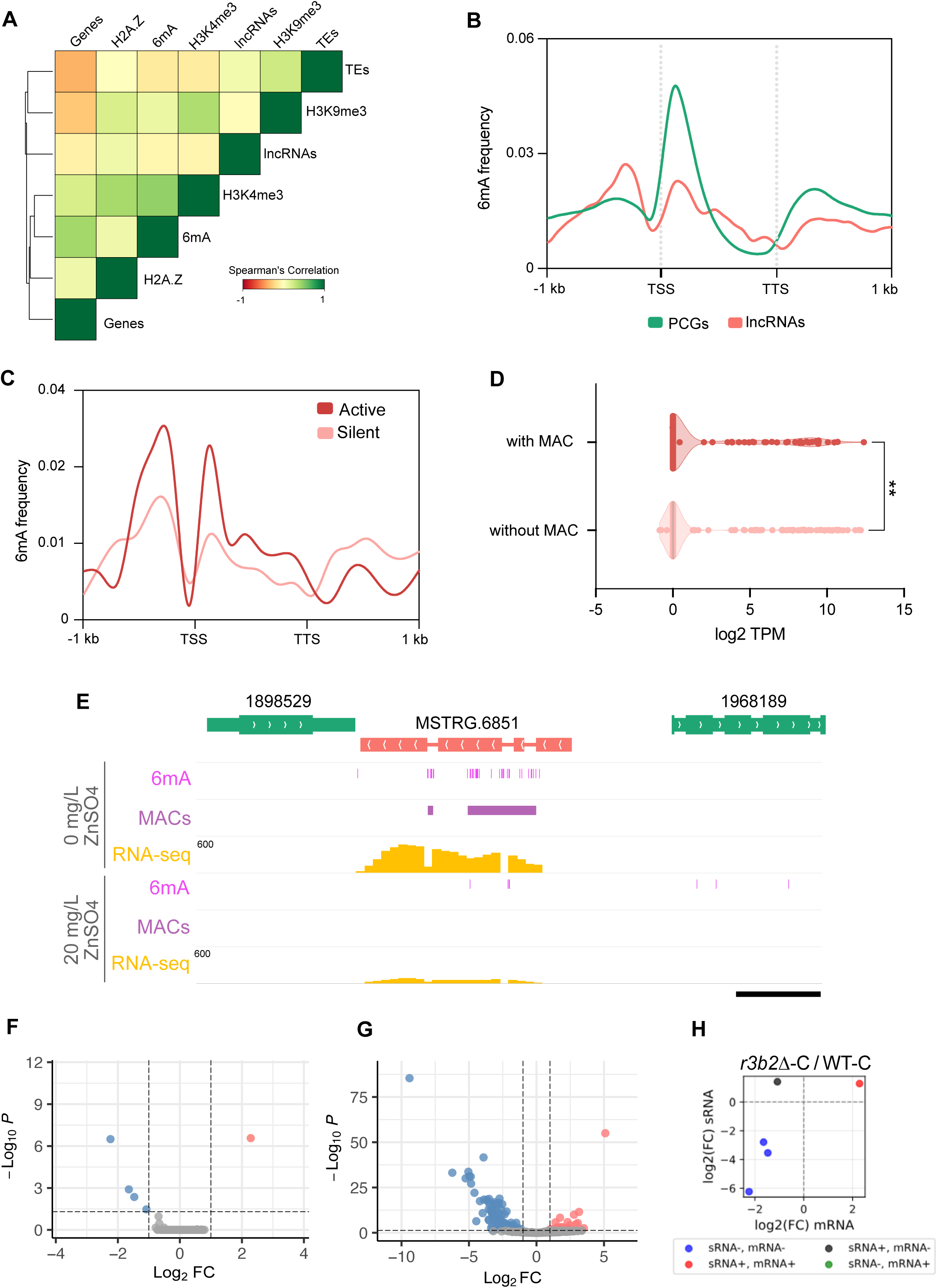
Epigenetic regulation of lncRNAs by methylation and by RNA interference. **(A)** Genomic distribution of lncRNAs relative to active (H3K4me3/6mA) and inactive (H3K9me3/transposon-rich) chromatin regions. lncRNAs show preferential enrichment in inactive regions. **(B)** Metagene analysis of 6mA distribution across lncRNAs compared with PCGs, showing a similar pattern with enrichment near promoters and downstream of TSS. **(C)** Frequency of 6mA in transcriptionally silent (TPM = 0) versus active lncRNAs, revealing higher methylation in active transcripts. **(D)** Expression levels of lncRNAs with or without a methylated adenine cluster (MAC) around the TSS (−300 to +300 bp). Methylated lncRNAs show significantly higher expression. **(E)** Example of lncRNA regulation by 6mA: loss of a MAC in MSTRG.5861 upon Mta1 knockdown correlates with reduced expression. **(F)** Volcano plots of the expression of lncRNAs in the *r3b2Δ* strain versus the WT strain interacting with macrophages (*r3b2Δ*-M/ WT-M). Red and blue dots indicate the upregulated and downregulated lncRNAs, respectively, while grey dots show not differentially expressed lncRNAs. **(G)** Volcano plots of the sRNA production from lncRNAs in the mutant versus the WT in non-stressful conditions (*r3b2Δ-*C / WT-C). **(H)** Scatter plot showing the relationship between the production of the sRNA and mRNA in the *r3b2Δ-*C / WT-C. Blue dots are lncRNAs with downregulation in their mRNA and sRNA, black dots are genes upregulated at the sRNA level and downregulation at the mRNA level, and the red dots are lncRNAs with simultaneous upregulation in the sRNA and mRNA.

### lncRNAs are targeted by canonical and non-canonical RNAi pathways in *M. lusitanicus*

The RNA interference (RNAi) machinery in *M. lusitanicus* includes two major pathways: the canonical RNAi pathway and the non-canonical RNAi pathway (NCRIP). The canonical pathway regulates gene expression and maintains genome stability^22^. In contrast, NCRIP is a known regulator of virulence, genome stability, and core developmental processes such as sporulation, largely by modulating the expression of genes involved in energy metabolism^15^. Given that lncRNAs often share regulatory patterns with protein-coding genes (PCGs), we examined whether lncRNAs are subject to NCRIP-mediated degradation by analyzing their small RNA (sRNA) signatures. Under non-stress conditions, comparison of the NCRIP-deficient mutant (*r3b2*Δ-C) and wild type (WT-C) revealed only five lncRNAs significantly differentially expressed at the mRNA level (**Figure 3F**). sRNA profiling of the same comparison uncovered 136 lncRNAs producing abundant sRNAs in the wild type (**Figure 3G**), consistent with active NCRIP-mediated degradation. Strikingly, the same five lncRNAs were simultaneously differentially expressed at both the mRNA and sRNA levels (**Figure 3H**), providing direct evidence of NCRIP-dependent regulation.

Interestingly, during phagocytosis of spores from the *r3b2*Δ strain, we detected 115 lncRNAs generating sRNAs, of which 38 showed coordinated changes at both the transcript and sRNA levels (**Supplementary** Figure 2A and 2B). This pattern indicates that these lncRNAs are targeted predominantly by the canonical RNAi pathway in the absence of NCRIP. Similarly, in wild-type spores confronted with macrophages, 36 lncRNAs produced a robust sRNA degradome, with 14 additionally showing differential mRNA expression (**Supplementary** Figure 2C and 2D), suggesting concurrent regulation by NCRIP and/or canonical RNAi.

### Specific lncRNAs display a high degree of evolutionary conservation in Mucorales

To assess the evolutionary conservation of lncRNAs in Mucorales, we performed a comparative genomic analysis including related species. Pairwise BLASTn searches were conducted using the lncRNA sequences from *M. lusitanicus* and *R. microsporus* against custom genomic databases of selected species. Hits overlapping annotated protein-coding genes or tRNAs (with ≥50% identity and ≥50% alignment length) were excluded to minimize false positives due to misannotated or misclassified sequences. The remaining matches were cross-referenced with unannotated intergenic regions, representing putative homologous lncRNAs (**Supplementary Data 3**). For *M. lusitanicus*, we analyzed the genomes of *R. microsporus*, *Ellisomyces anomalus*, *Mucor ambiguus*, *Mucor circinelloides*, *Parasitella parasitica*, and *Mucor racemosus* (**Figure 4A**). For *Rhizopus microsporus*, we considered five closely related taxa: *R. microsporus var. chinensis*, *Amylomyces rouxii*, *Rhizopus delemar*, *Rhizopus stolonifer*, and *Sporodiniella umbellata* and well as *M. lusitanicus* (**Figure 4B**). The disparity in the degree of conservation led us to focus our analysis on those lncRNAs that were conserved across at least five or three of the compared species for *M. lustianicus* and *R. microsporus*, respectively. Despite the expected low sequence conservation of fungal lncRNAs, a subset of lncRNAs showed evidence of conservation across multiple species. In *M. lusitanicus*, six lncRNAs exhibited conserved sequence similarity with unannotated regions in seven different species, including *R. microsporus* (**Figure 4A, red line**). Moreover, an additional group of 42 lncRNAs was found to be conserved across four *Mucor* species (**Figure 4A, blue line**). Remarkably, a single *R. microsporus* lncRNA (MSTRG.4826) aligned with unannotated genomic regions in seven Mucorales genomes, including *M. lusitanicus*, suggesting a potential ancestral origin (**Figure 4B**). Interestingly, its ortholog in *M. lusitanicus*, MSTRG.5947, is conserved across all examined *Mucor* species. Additionally, the *R. microsporus* lncRNA MSTRG.3902 was conserved across six related species, including *M. lusitanicus* (**Figure 4B**). Finally, the lncRNA MSTRG.4436 represents a third case of conservation between *M. lusitanicus* and *R. microsporus*.

**Fig. 4.**
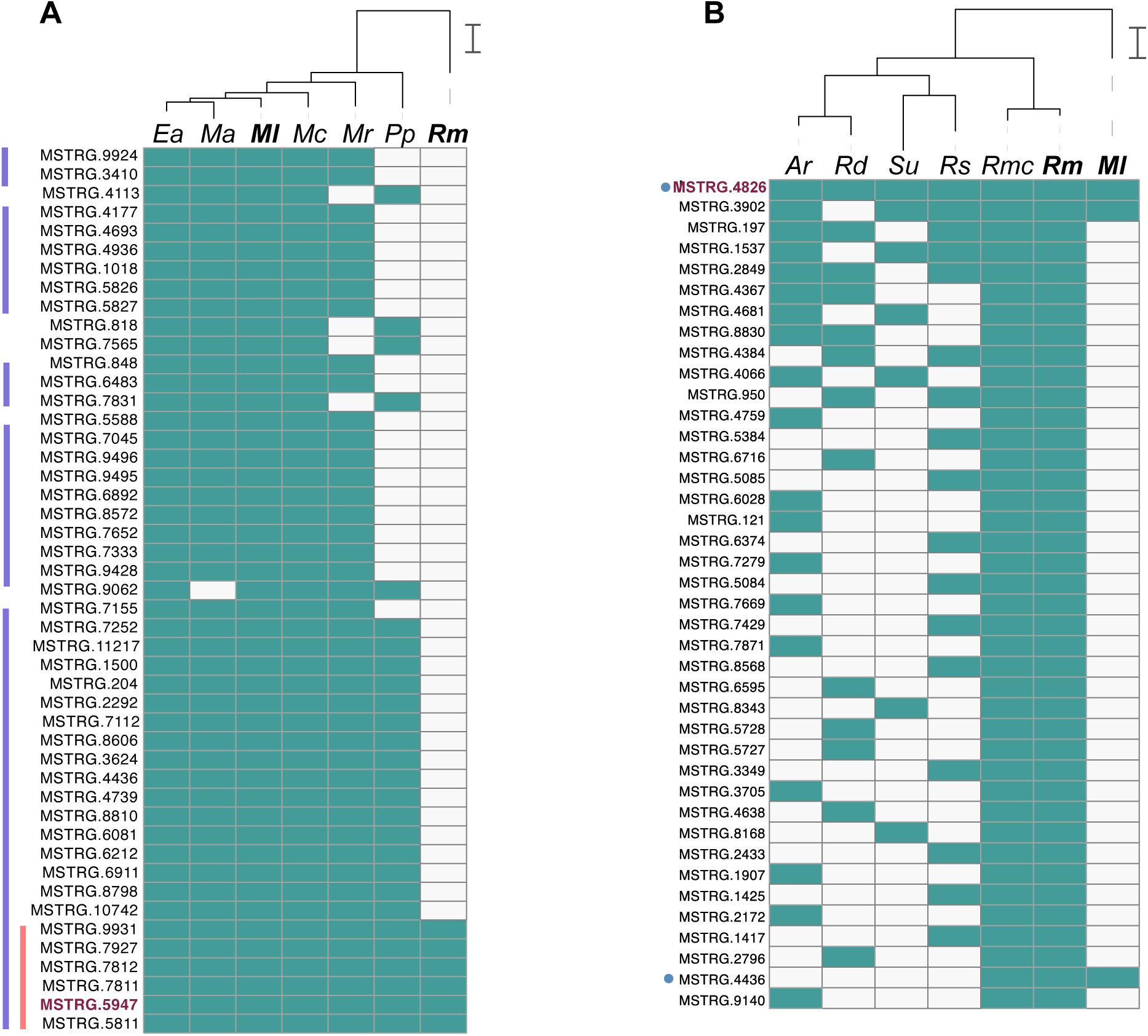
Conservation of lncRNAs across Mucorales species. Comparative genomic analysis of lncRNAs from *Mucor lusitanicus* (Ml) and *Rhizopus microsporus* (Rm) across ten closely related Mucorales genomes: *Ellisomyces anomalus* (Ea), *Mucor ambiguus* (Ma), *Mucor circinelloides* (Mc), *Parasitella parasitica* (Pp), *Mucor racemosus* (Mr), *R. microsporus* var. *chinensis* (Rmc), *Amylomyces rouxii* (Ar), *R. delemar* (Rd), *R. stolonifer* (Rs), and *Sporodiniella umbellata* (Su). (A) Number of *M. lusitanicus* lncRNAs conserved in at least five species. Six lncRNAs (red line) are conserved in seven species, including *R. microsporus*. Blue lines indicate lncRNAs conserved across all *Mucor* species (Ml, Ea, Ma, Mc, Mr). (C) Number of *R. microsporus* lncRNAs conserved in at least three related species. One lncRNA (blue dot), MSTRG.4826 (orthologous to *M. lusitanicus* MSTRG.5947), is conserved in all analyzed genomes, indicating deep evolutionary conservation. Another, MSTRG.4436, is conserved in *M. lusitanicus*, *R. microsporus,* and *R. microsporus* var. *chinensis*.

### Role of lncRNAs in the pathogenesis of Mucorales

Expression data of *M. lusitanicus* WT and NCRIP-deficient *r3b2*Δ mutant strain confronted with macrophages was evaluated to assess the extent of lncRNAs implication in this response. 212 different lncRNAs were differentially expressed in the confrontation of the WT strain versus a non-infectious context (**Supplementary** Figure 3A), and 209 were differentially expressed in the interaction of the *r3b2*Δ mutant compared to saprophytic growth (**Supplementary** Figure 3B). 163 genes, representing 63% of the total DE lncRNAs (DELs), are commonly differently expressed in the two comparisons (**Supplementary** Figure 3C). These include lncRNAs that are significantly upregulated or downregulated in both wild-type and mutant strains upon exposure to phagocytic cells. This implication of lncRNAs in infection-related processes was not exclusive to *M. lusitanicus*. In the case of *R. microsporus,* we performed a differential expression analysis using the RNA-seq data derived from an infected murine model. When focusing on the identified lncRNAs, we found a total of 129 lncRNAs displaying significant differential expression during infection (adjusted *p* < 0.05, |log2FC| > 1), including 38 upregulated and 91 downregulated lncRNAs, among them MSTRG.4436, MSTRG.1050, and MSTRG.7756 (**Figure 5A**). In addition, we took advantage of the dataset to evaluate the involvement of the lncRNAs characterized in mice in the response to Mucorales infection. A total of 881 host lncRNAs showed significant expression changes, with 424 upregulated and 457 downregulated during the interaction with *R. microsporus* (**Figure 5B**). This robust transcriptional response reflects widespread modulation of the host non-coding transcriptome in response to fungal challenge.

**Fig. 5.**
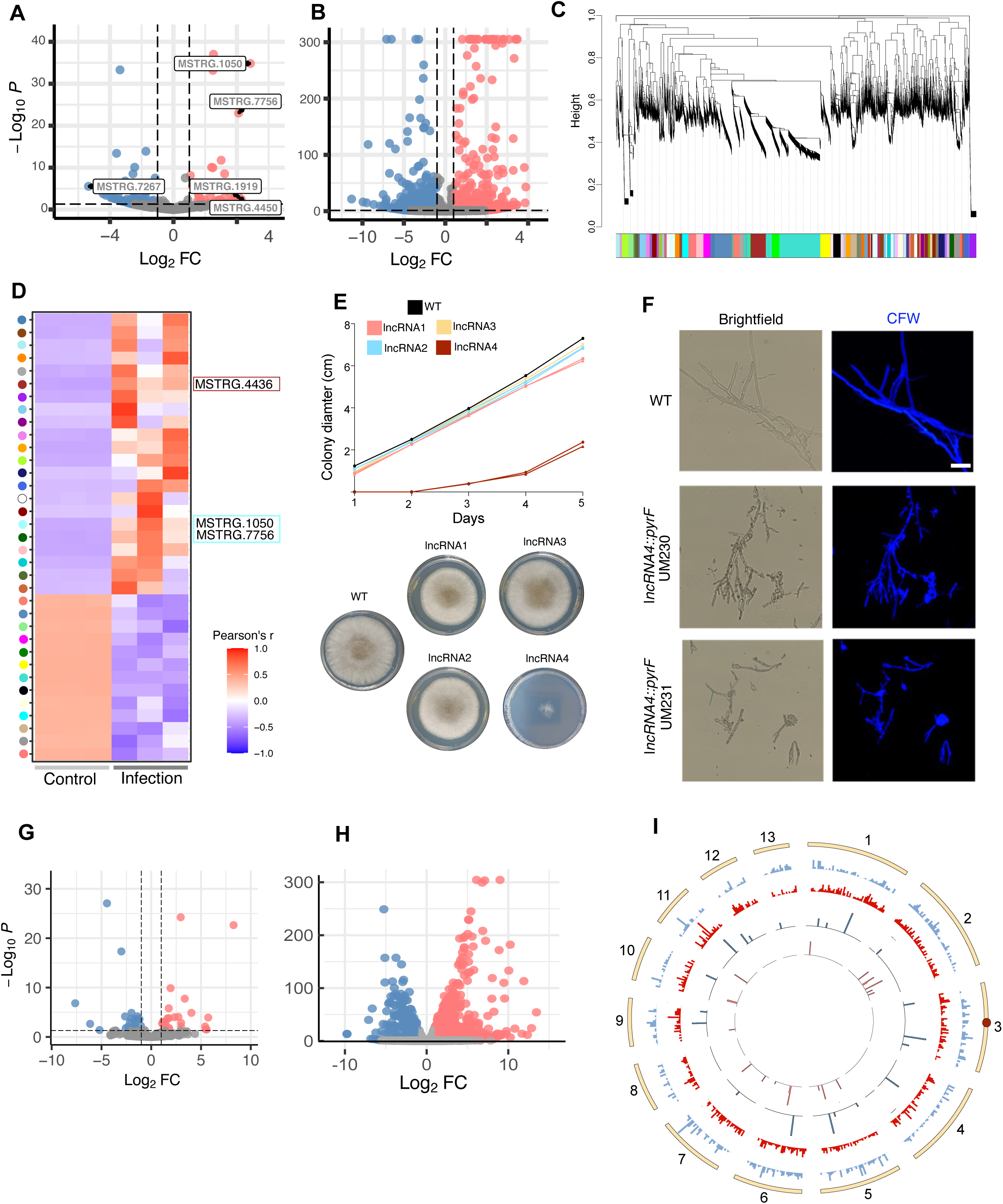
Differential expression, co-expression network analysis, and functional characterization of lncRNAs in *R. microsporus*. **(A**) Volcano plot of differentially expressed lncRNAs in *R. microsporus* under infection vs saprophytic growth; the most significantly upregulated and downregulated lncRNAs are highlighted. **(B)** Volcano plot of lncRNA expression in infected versus non-infected mouse tissue. **(C)** Weighted gene co-expression network analysis (WGCNA) of the differentially expressed genes and lncRNAs of *R. microsporus* under saprophytic and infection conditions. Dendrogram of clustered genes based on topological overlap, with modules indicated by distinct colors. **(D)** Heatmap of module–trait relationships showing Pearson correlation coefficients between module eigengenes and infection status. Blue, negative correlation; red, positive correlation. Three lncRNAs (MSTRG.4436, MSTRG.1050, MSTRG.7756) are assigned to modules strongly associated with infection. **(E)** Radial growth of wild type (WT) and lncRNA mutants over 5 days (mean ± s.d., n = 3). Mutants lacking lncRNA1 and lncRNA4 showed reduced growth, with lncRNA4⁻ exhibiting the strongest effect. **(F)** Hyphal morphology of WT and lncRNA4⁻ mutants. Mutants display shorter, less branched filaments. Scale bars, 10 μm. **(G)** Volcano plot of lncRNA expression in lncRNA4⁻ versus WT strains. Red, upregulated; blue, downregulated; grey, not differentially expressed. **(H)** Differential expression of protein-coding genes (PCGs) in lncRNA4⁻ versus WT. **(I)** Circos plot of differentially expressed PCGs and lncRNAs across all scaffolds of the *R. microsporus* genome. Tracks (outer to inner): upregulated PCGs (blue), downregulated PCGs (red), upregulated lncRNAs (blue), downregulated lncRNAs (red). The lncRNA4⁻ locus on scaffold 3 is highlighted. Gene density was computed in 50 kb windows.

To explore potential functional roles of these lncRNAs, a weighted gene co-expression network was constructed using WGCNA, incorporating both differentially expressed lncRNAs and protein-coding genes (PCGs). Hierarchical clustering based on the topological overlap matrix (TOM) identified 35 distinct co-expression modules, each assigned a unique color (**Figure 5C**). Module eigengenes (MEs) were calculated to summarize module expression patterns, and their correlations with infection status were visualized as a heatmap (**Figure 5D**). Several modules displayed strong positive or negative correlations with infection, indicating that the co-expressed lncRNAs and PCGs within these modules may play functional roles during pathogenesis. Among the positively correlated modules, we identified individual modules containing the lncRNAs MSTRG.4436, MSTRG.1050, and MSTRG.7756, with each lncRNA located in a separate module.

### Targeted disruption of infection-induced and conserved lncRNAs reveals their essential roles in *Rhizopus microsporus*

To further explore the biological implications of identified lncRNAs, we generated CRISPR/Cas9-assisted knock-out mutant strain in four selected candidates: lncRNA1 (MSTRG.4436), lncRNA2 (MSTRG.1050), lncRNA3 (MSTRG.7756) based on their significant upregulation during murine infection as well as lncRNA4 (MSTRG.4826), which was found to exhibit broad evolutionary conservation in Mucorales species analyzed (**Supplementary** Figure 4). Noteworthy, independently of the number of growth cycles in selective media, we were unable to isolate homokaryotic mutants for lncRNA2 and lncRNA4, suggesting that they may be essential for *R. microsoporus* survival (**Supplementary** Figure 4). Phenotypic characterization revealed that the heterokaryotic lncRNA2 mutant and the lncRNA3 deletion mutant did not display significant growth alterations (Figure 5E). In contrast, the lncRNA1 mutant exhibited a modest reduction in growth rate (p = 0.0012, Welch’s t-test), while the heterokaryotic lncRNA4 mutant showed a markedly pronounced growth defect (p < 0.0001, Welch’s t-test) (Figure 5E). Moreover, further inspection of the lncRNA4 defective mutant strains revealed a clear defective filamentous growth in which hyphae appeared shorter and truncated compared to the WT strain (**Figure 5F**). The essentiality of the identified lncRNA2 and lncRNA4 prompted us to further dissect the regulatory implications of these two lncRNAs in the biology of *R. microsporus*. Transcriptomic analyses of both mutants uncovered candidate target genes and pathways that may underlie their critical roles in fungal growth and survival. In the lncRNA4⁻ mutant, 48 DELs and 1607 differentially expressed protein-coding genes (PCGs) were identified (padj < 0.05, |log₂FC| ≥ 1). DELs comprised 25 upregulated and 23 downregulated transcripts (**Figure 5G**), whereas PCGs included 1037 upregulated and 570 downregulated genes (**Figure 5H**). GO enrichment analyses highlighted strong overrepresentation of oxidoreductase and hydrolase activities, particularly those linked to carbohydrate metabolism and cell wall remodeling (e.g., chitinase, chitin deacetylase). Additional enrichments were detected in amino acid metabolism, organic acid catabolism, and transmembrane transport of carbohydrates and metal ions (**Supplementary** Figure 5A**, right panel**). These findings indicate that lncRNA4 orchestrates redox homeostasis, nutrient transport, and cell wall integrity, processes essential for fungal viability. Genomic distribution analyses further revealed that lncRNA4 putative targets are dispersed throughout the genome (**Figure 5I**).

In contrast, the lncRNA2 mutant exhibited 31 DELs and 876 differentially expressed PCGs (**Supplementary** Figure 5B and 5C). DELs comprised 13 upregulated and 18 downregulated transcripts, while PCGs included 405 upregulated and 471 downregulated genes. GO enrichment revealed a striking dominance of mitochondrial processes, including protein targeting and insertion into the inner mitochondrial membrane, intermembrane space chaperone complexes, and overall mitochondrial organization. Ribosomal components and structural constituents of the ribosome were also significantly enriched, pointing to altered protein synthesis. Additional functional categories included oxidoreductase activity, heme and tetrapyrrole binding, zinc ion transport, arginine metabolism, and glutathione transferase activity, consistent with roles in redox balance and metabolic stress responses (**Supplementary** Figure 5A**, left panel**).

## DISCUSSION

This study represents the first extensive characterization and functional analysis of lncRNAs in early-diverging fungi (EDF), focusing primarily on *M. lusitanicus* and *R. microsporus*, two representative species of this group with biological and clinical importance. Our findings contribute significantly to the understanding of fungal lncRNA biology, their evolutionary conservation, regulation, and crucial roles in fungal pathogenicity. While lncRNAs have been widely studied in plants^23,24^, animals^25^, and some fungal groups^9,14,16,17^, their presence, characteristics, and functions in early-diverging fungi have remained largely unexplored. Consistent with observations in other fungi^13,14,16,17^, lncRNAs in *M. lusitanicus* and *R. microsporus* display canonical eukaryotic features, including shorter transcript lengths, lower GC content, and reduced expression relative to protein-coding genes (PCGs), reinforcing their classification as bona fide lncRNAs. Furthermore, the abundance of lncRNAs correlated with local PCG density, suggesting that their transcription is influenced by genome organization and chromatin context. Comparative analysis revealed species-specific lncRNA repertoires, reflecting both genome architecture and evolutionary divergence, with enrichment in the inactive compartment of chromatin but with evidence of regulation with 6mA, a rare epigenetic characteristic of this particular group of fungi^21^.

In addition to epigenetic regulation, our data demonstrate that lncRNAs are subject to post-transcriptional control via RNA interference in *M. lusitanicus*. The non-canonical RNAi pathway (NCRIP) targets many lncRNAs, as revealed by sRNA degradome analysis, with over a hundred loci undergoing active degradation in the wild type despite few changes at the mRNA level under non-stress conditions. During macrophage interaction, some lncRNAs are regulated by both NCRIP and canonical RNAi, indicating a coordinated RNAi-mediated control responsive to environmental and host cues. These findings are consistent with previous observations in filamentous fungi, where RNAi modulates both coding and non-coding transcripts to support stress adaptation and genome stability^26,27^.

Together, these results suggest that lncRNAs in Mucorales are embedded within multilayered regulatory networks: transcription is influenced by chromatin context and 6mA-mediated epigenetic signals, while transcript stability is actively controlled by RNAi pathways. This complex regulatory interplay positions lncRNAs as modulators or scaffolds within gene expression networks relevant to host-pathogen interactions, virulence, and stress adaptation, highlighting their potential as critical regulators of fungal growth and as targets for therapeutic intervention.

Our comparative genomics analysis revealed that the majority of Mucorales lncRNAs display low primary sequence conservation, consistent with rapid evolutionary turnover and species-specific emergence^28,29^. Nonetheless, a subset of lncRNAs exhibited significant sequence conservation across multiple species, suggesting potential functional relevance. Notably, we identified one lncRNA conserved across eight Mucorales genomes, including *M. lusitanicus* and *R. microsporus*, implying a critical biological role potentially linked to conserved regulatory networks or fundamental aspects of fungal physiology. These observations suggest that while most lncRNAs are species-specific, a conserved minority may perform essential functions, warranting further functional characterization and structural analysis. Expression studies revealed that lncRNAs in *R. microsporus* and *M. lusitanicus*, similar to PCGs^30^, are dynamically regulated during murine and macrophage infections, respectively, underscoring their potential roles in host-pathogen interactions. A subset of fungal lncRNAs was significantly induced *in vivo*, while host lncRNAs also exhibited broad transcriptional changes, reflecting extensive modulation of the host non-coding transcriptome in response to fungal challenge. These findings align with observations in *Candida albicans*, where lncRNAs were modulated during epithelial infection^9^, suggesting functional relevance in fungal pathogenesis. Similarly, in *C. neoformans*, lncRNAs display dynamic expression patterns during murine macrophage infection^31^, further supporting their involvement in host-pathogen interactions.

Targeted disruption of infection-induced lncRNAs demonstrated that certain loci, particularly lncRNA4 and lncRNA2, are essential for fungal viability. LncRNA4 mutants displayed severe growth defects, abnormal hyphal morphology, and nuclear heterogeneity, with transcriptomic analyses indicating roles in redox homeostasis, nutrient transport, and cell wall remodeling. LncRNA2 predominantly regulated mitochondrial function, protein synthesis, and redox balance.

These results not only reveal fundamental aspects of lncRNA biology in early-diverging fungi but also highlight potential targets for antifungal strategies. By integrating chromatin state, epigenetic modifications, and RNAi-dependent turnover, lncRNAs emerge as critical regulatory hubs that contribute to fungal adaptation, pathogenicity, and survival under host-imposed stress. Future studies dissecting the mechanistic links between 6mA deposition, RNAi-mediated degradation, and lncRNA function will be essential to fully understand their contribution to fungal biology and disease.

In summary, our work establishes lncRNAs as integral components of the Mucorales transcriptome, showing conserved evolutionary signatures, dynamic regulation during host infection, and essential roles in fungal viability and pathogenicity. Genetic and phenotypic analyses demonstrate for the first time that specific lncRNAs are indispensable for *R. microsporus* survival, identifying them as potential targets for antifungal intervention. Future studies dissecting their mechanistic roles, interacting partners, and downstream regulatory pathways will be crucial for a comprehensive understanding of fungal biology and infection. Ultimately, these insights may enable innovative antifungal therapies aimed at disrupting lncRNA-dependent regulatory networks, addressing the urgent clinical challenges posed by mucormycosis and related infections.

## ACKNOWLEDGMENTS

This research was supported by the grant PGC2018-097452-B-I00, PID2021-124674NB-I00, and PID2024-160088NB-I00 to F.E.N. and V.G, funded by MICIU/AEI/10.13039/501100011033 and by ERDF/EU. TG group acknowledges support from the Spanish Ministry of Science and Innovation (grant numbers PID2021-126067NB-I00, CPP2021-008552, PCI2022-135066-2, PLEC2023-010225, and PDC2022-133266-I00), cofounded by ERDF “A way of making Europe”, as well as support from the Catalan Research Agency (AGAUR) (grant number SGR01551); “La Caixa” foundation (grant number LCF/PR/HR21/00737), and Instituto de Salud Carlos III (CIBERINFEC CB21/13/00061-ISCIII-SGEFI/ERDF).

## AUTHOR CONTRIBUTIONS

G.T. performed the majority of the experimental work and bioinformatic formal analyses, curated data, performed visualization by creating figures and tables, and wrote and edited the original draft with substantial input from C.L., H.H., F.E.N., T.G., and V.G. H.H. contributed to formal bioinformatic analyses for lncRNA identification. C.L. conceptualized and conducted experimental work in mutant generation, performed formal analyses with bioinformatic data and performed visualization by creating figures and tables. E.N. provided resources. F.E.N., T.G., and V.G. acquired funds, administered the project and supervised the research and reviewed and edited the original draft. G.T. and V.G. conceptualized the project

## DATA AVAILABILITY

The raw sequence data that support the findings of this study have been deposited in the Gene Expression Omnibus (GEO) under accession number GSE308583.

## CODE AVAILABILITY

The scripts used for the bioinformatic analysis are available at https://github.com/ghizlanetahiri95/Mucorales_lncRNAs.

**Fig. Supp. 1 Distribution of lncRNAs and PCGs per scaffold (macro-scale) in *M. lusitanicus* (A) and *R. microsporus* (B).** Lines indicate the total number of genes per scaffold for each type, with all scaffolds from each genome assembly represented. PCGs are shown in green and lncRNAs in red, revealing similar scaffold-level distribution patterns and highlighting scaffolds with high gene content for both categories.

**Fig. Supp. 2 Regulation of lncRNAs by macrophage confrontation and RNAi pathways in *M. lusitanicus*.** Degradome analysis of sRNA–mRNA correspondence**. (A)** Volcano plot of sRNA production from lncRNAs in the *r3b2Δ* mutant with (M) and without (C) macrophages**. (B)** Scatter plot of sRNA versus mRNA production in *r3b2Δ* (C/M). Blue, lncRNAs with downregulation of both sRNA and mRNA; black, lncRNAs with sRNA upregulation and mRNA downregulation; red, lncRNAs with simultaneous sRNA and mRNA upregulation. **(C)** Volcano plot of sRNA production from lncRNAs in the WT strain with (M) and without (C) macrophages. **(D)** Scatter plot of sRNA versus mRNA production in WT (M/C), color-coded as in (B).

**Fig. Supp. 3 Regulation of lncRNAs by phagocytosis and RNAi pathways in *M. lusitanicus*. (A)** Volcano plot of differentially expressed lncRNAs in the WT strain with (M) and without (C) macrophages. **(B)** Volcano plot of differentially expressed lncRNAs in the *r3b2Δ* mutant during macrophage confrontation. **(C)** Venn diagram of lncRNAs regulated by both phagocytosis and RNAi pathways.

**Fig. Supp. 4 PCR validation of *R. microsporus* lncRNA disruption strains.** PCRs were performed to assess homokaryosis of lncRNA candidate mutants using primers ∼1 kb outside the 38 bp homology regions. Amplification products for the disrupted lncRNAs (red arrows) were 4.2 kb for lncRNA1–3 and 4.3 kb for lncRNA4; WT versions yielded 0.7 kb (lncRNA1–3) and 0.8 kb (lncRNA4) fragments. Red rows indicate disrupted lncRNAs, black rows indicate WT versions. Mutants of lncRNA1 and lncRNA3 were homokaryotic, whereas lncRNA2 and lncRNA4 remained heterokaryotic after multiple vegetative cycles.

**Fig. Supp. 5 Differential expression and functional enrichment analyses in lncRNA mutants. (A)** Gene Ontology (GO) enrichment analysis of differentially expressed genes (DEGs) in the lncRNA4⁻ mutant (right) and lncRNA2⁻ mutant (left). Circle size represents the gene ratio; color scale indicates –log(p-value). **(B)** Volcano plot of lncRNAs differentially expressed in the lncRNA2⁻ mutant relative to WT. **(C)** Volcano plot of protein-coding genes (PCGs) differentially expressed in the lncRNA2⁻ mutant relative to WT. Differential expression and functional enrichment analyses in lncRNA mutants.

