## Supplementary information for "Genome-wide Discovery of lncRNAs in Mucorales Reveals Essential Roles in Development and Fungal Biology"

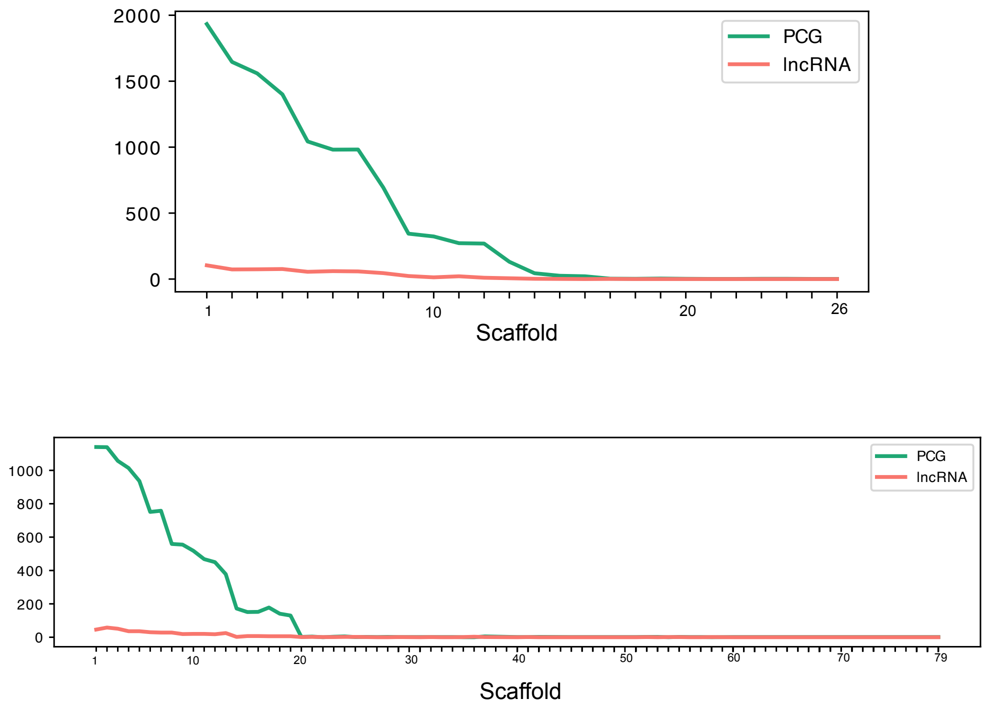


**Fig. Supp. 1 Distribution of lncRNAs and PCGs per scaffold (macro-scale) in *M. lusitanicus* (A) and *R. microsporus* (B).** Lines indicate the total number of genes per scaffold for each type, with all scaffolds from each genome assembly represented. PCGs are shown in green and lncRNAs in red, revealing similar scaffold-level distribution patterns and highlighting scaffolds with high gene content for both categories.

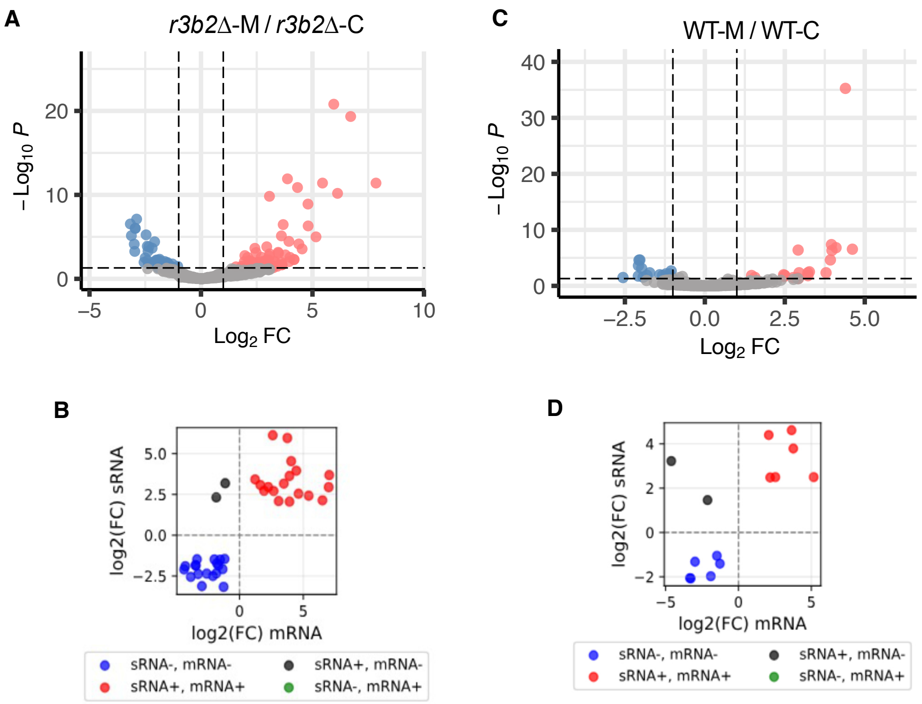


**Fig. Supp. 2 Regulation of lncRNAs by macrophage confrontation and RNAi pathways in *M. lusitanicus*.** Degradome analysis of sRNA–mRNA correspondence**. (A)** Volcano plot of sRNA production from lncRNAs in the *r3b2Δ* mutant with (M) and without (C) macrophages**. (B)** Scatter plot of sRNA versus mRNA production in *r3b2Δ* (C/M). Blue, lncRNAs with downregulation of both sRNA and mRNA; black, lncRNAs with sRNA upregulation and mRNA downregulation; red, lncRNAs with simultaneous sRNA and mRNA upregulation. **(C)** Volcano plot of sRNA production from lncRNAs in the WT strain with (M) and without (C) macrophages. **(D)** Scatter plot of sRNA versus mRNA production in WT (M/C), color-coded as in (B).

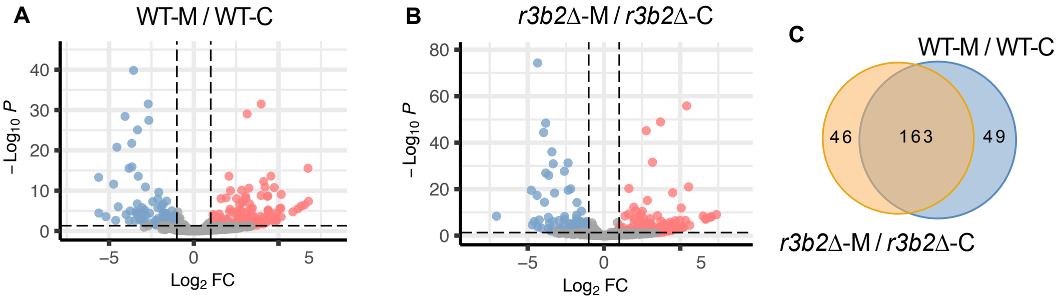


**Fig. Supp. 3 Regulation of lncRNAs by phagocytosis and RNAi pathways in *M. lusitanicus*. (A)** Volcano plot of differentially expressed lncRNAs in the WT strain with (M) and without (C) macrophages. **(B)** Volcano plot of differentially expressed lncRNAs in the *r3b2Δ* mutant during macrophage confrontation. **(C)** Venn diagram of lncRNAs regulated by both phagocytosis and RNAi pathways.

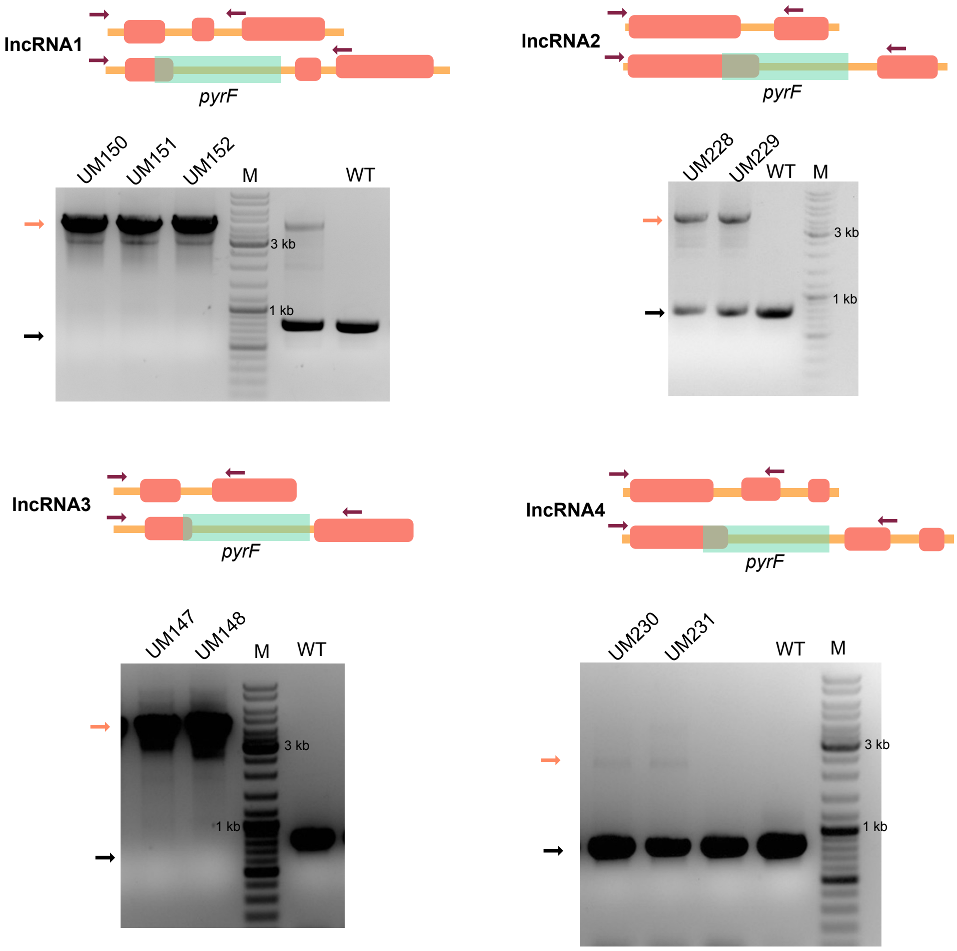


**Fig. Supp. 4 PCR validation of *R. microsporus* lncRNA disruption strains.** PCRs were performed to assess homokaryosis of lncRNA candidate mutants using primers ~1 kb outside the 38 bp homology regions. Amplification products for the disrupted lncRNAs (red arrows) were 4.2 kb for lncRNA1–3 and 4.3 kb for lncRNA4; WT versions yielded 0.7 kb (lncRNA1–3) and 0.8 kb (lncRNA4) fragments. Red rows indicate disrupted lncRNAs, black rows indicate WT versions. Mutants of lncRNA1 and lncRNA3 were homokaryotic, whereas lncRNA2 and lncRNA4 remained heterokaryotic after multiple vegetative cycles.

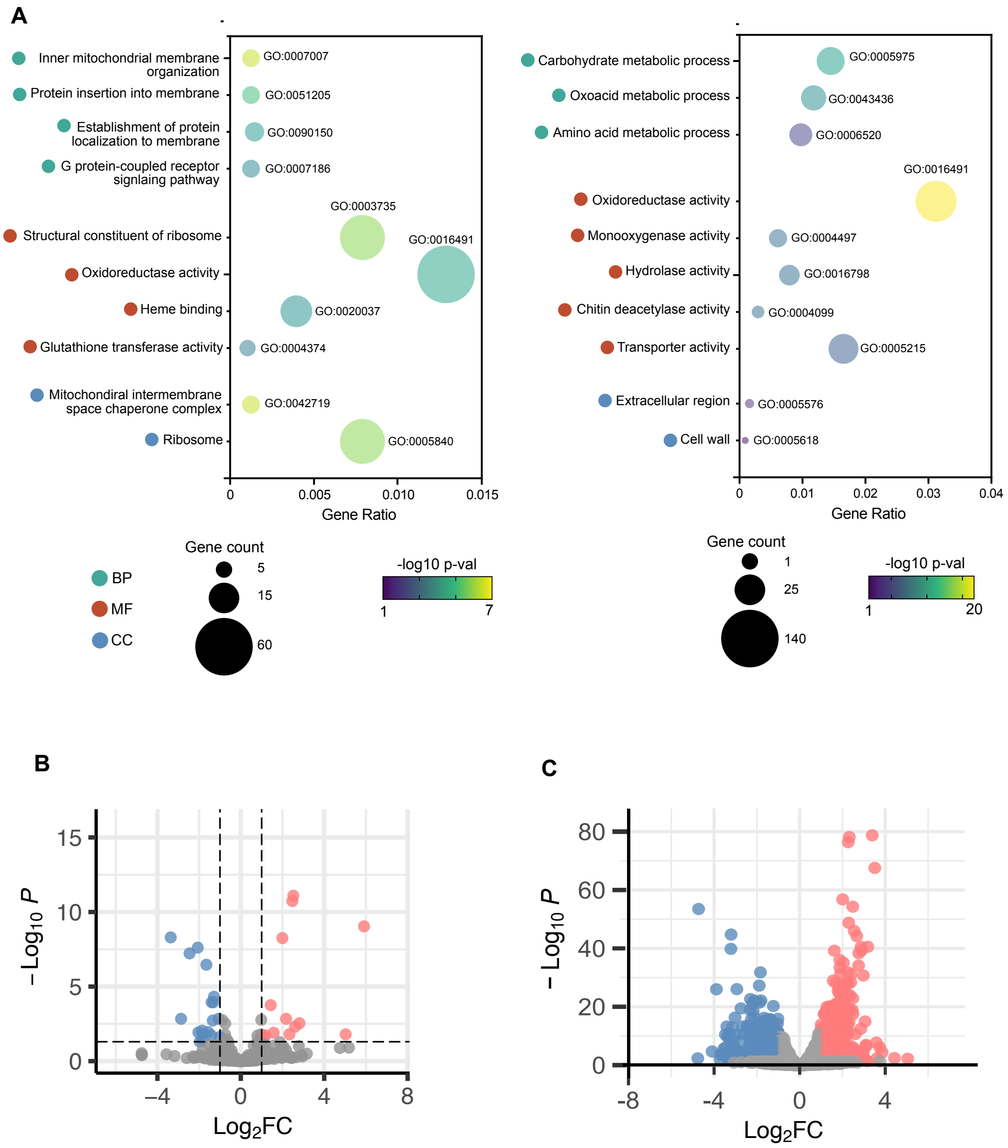


**Fig. Supp. 5 Differential expression and functional enrichment analyses in lncRNA mutants. (A)** Gene Ontology (GO) enrichment analysis of differentially expressed genes (DEGs) in the lncRNA4⁻ mutant (right) and lncRNA2⁻ mutant (left). Circle size represents the gene ratio; color scale indicates –log(p-value). **(B)** Volcano plot of lncRNAs differentially expressed in the lncRNA2⁻ mutant relative to WT. **(C)** Volcano plot of protein-coding genes (PCGs) differentially expressed in the lncRNA2⁻ mutant relative to WT. Differential expression and functional enrichment analyses in lncRNA mutants.

**Supplementary Table 1**

| Strain | Genotype | Description | Organism | Source |
|---|---|---|---|---|
| ATCC11559 | WT | WT strain | *R. microsporus* | Lax et al. 2021 |
| UM33 | *pyrF-, leuA-* | Avirulent Control. Used as a receptor strain for the mutant generation | *R. microsporus* | Tahiri et al. 2025 |
| UM6 | *pyrF+, leuA-* | Virulent strain | *R. microsporus* | Lax et al. 2021 |
| UM150 | *leuA- ,lncRNA1::pyrF* | Mutant in which the lncRNA1 has been disrupted with the *pyrF* gene. Derived from UM33 | *R. microsporus* | This work |
| UM151 | *leuA-, lncRNA1::pyrF* | Mutant in which the lncRNA1 has been disrupted with the *pyrF* gene. Derived from UM33 | *R. microsporus* | This work |
| UM152 | *leuA-, lncRNA1::pyrF* | Mutant in which lncRNA1 has been disrupted with the *pyrF* gene. Derived from UM33 | *R. microsporus* | This work |
| UM228 | *leuA-, lncRNA2::pyrF* | Mutant in which the lncRNA2 has been disrupted with the *pyrF* gene. Derived from UM33 | *R. microsporus* | This work |
| UM229 | *leuA-, lncRNA2::pyrF* | Mutant in which the lncRNA2 has been disrupted with the pyrF gene. Derived from UM33 | *R. microsporus* | This work |
| UM147 | *leuA-, lncRNA3::pyrF* | Mutant in which the lncRNA3 has been disrupted with the *pyrF* gene. Derived from UM33 | *R. microsporus* | This work |
| UM148 | *leuA-, lncRNA3::pyrF* | Mutant in which the lncRNA3 has been disrupted with the *pyrF* gene. Derived from UM33 | *R. microsporus* | This work |
| UM230 | *leuA-, lncRNA4::pyrF* | Mutant in which the lncRNA4 has been disrupted with the *pyrF* gene. Derived from UM33 | *R. microsporus* | This work |
| UM231 | *leuA-, lncRNA4::pyrF* | Mutant in which the lncRNA4 has been disrupted with the *pyrF* gene. Derived from UM33 | *R. microsporus* | This work |

**Supplementary Table 2**

| | Target gene | 5'3' |
|---|---|---|
| gRNAs for lncRNA mutants in *R. microsporus* | gRNA_lncRNA1 | CATCAGGTCTTCTTTCACTG |
| | gRNA_lncRNA2 | AAAGAGAACTCAAACAAAGC |
| | gRNA_lncRNA3 | CTGTCCGCGACATCGTTTGT |
| | gRNA_lncRNA4 | AGTGTTTGCCCAACACTGAG |

**Supplementary Table 3**

| | Amplified gene | 5'3' |
|---|---|---|
| lncRNA mutants in *R. microsporus* | pyrF_F_lncRNA1_Fw | ATTACTGTATGGTACGCTTTGCATCAGGTCTTCTTTCATCCTCCATAAGAATTTGACAG |
| | pyrF_F_lncRNA1_Rv | TTTTTTACATTCAAGAGTTATACTTACATAAACCTCAGTGATAAAACGAAGATGTGGCTGTC |
| | pyrF_F_lncRNA2_Fw | CAGCAAATGCTTAAATACGTAAAAGAGAACTCAAACAATCCTCCATAAGAATTTGACAG |
| | pyrF_F_lncRNA2_Rv | CAATTTTGGCTGACTAACTGATAATGTGCTAGCCAGCTTGATAAAACGAAGATGTGGCTGTC |
| | pyrF_F_lncRNA3_Fw | GGCTAGCGATATCCATTTTGACTGTCCGCGACATCGTTTCCTCCATAAGAATTTGACAG |
| | pyrF_F_lncRNA3_Rv | TGTGCTTTATTTATTTTTCAAATACAGTGCTTCCAACATGATAAAACGAAGATGTGGCTGTC |
| | pyrF_F_lncRNA4_Fw | ATTCCATCAGACATCCAAGGAAGTGTTTGCCCAACACTTCCTCCATAAGAATTTGACAG |
| | pyrF_F_lncRNA4_Rv | GTTTAGTAGGAGCACTTTGTGCTCCGAAAGATCCTCTCTGATAAAACGAAGATGTGGCTGTC |
| | lncRNA1_locus_Fw | TAACAAAGTAAGTATTATCTAC |
| | lncRNA1_locus_Rv | GTTACTAAAATCAATATATTAC |
| | lncRNA2_locus_Fw | CTTTATGATGCTTGAGGATGTG |
| | lncRNA2_locus_Rv | ACACACATTTCTATCCCAATGC |
| | lncRNA3_locus_Fw | TGACATGGTTTCTAAAAATACG |
| | lncRNA3_locus_Rv | TGGCACTCGACAATATAAACAG |
| | lncRNA4_locus_Fw | AATGCATACACACACTTAAAG |
| | lncRNA4_locus_Rv | CGAGCCACTGCCTCAGTTTAAC |
